# The pyruvate dehydrogenase complex regulates matrix protein phosphorylation and mitophagic selectivity

**DOI:** 10.1101/2022.03.16.484611

**Authors:** Panagiota Kolitsida, Vladimir Nolic, Jianwen Zhou, Michael Stumpe, Natalie M. Niemi, Jörn Dengjel, Hagai Abeliovich

## Abstract

The mitophagic degradation of mitochondrial matrix proteins in *S. cerevisiae* was previously shown to be selective, reflecting a pre-engulfment sorting step within the mitochondrial network. This selectivity is regulated through phosphorylation of mitochondrial matrix proteins by the matrix kinases Pkp1 and Pkp2, which in turn appear to be regulated by the phosphatase Aup1/Ptc6. However, these same proteins also regulate the phosphorylation status and catalytic activity of the yeast pyruvate dehydrogenase complex, which is critical for mitochondrial metabolism. To understand the relationship between these two functions, we evaluated the role of the pyruvate dehydrogenase complex in mitophagic selectivity. Surprisingly, we identified a novel function of the complex in regulating mitophagic selectivity, which is independent of its enzymatic activity. Our data support a model in which the pyruvate dehydrogenase complex directly regulates the activity of its associated kinases and phosphatases. This regulatory interaction then determines the phosphorylation state of mitochondrial matrix proteins and their mitophagic fates.

## Introduction

Mitochondrial autophagy, or mitophagy, is a quality control mechanism that culls defective mitochondria in an effort to prevent cellular damage and dysfunction. Indeed, defects in mitophagic mechanisms have been shown to underlie human pathologies that can be traced to mitochondrial dysfunction. These include Parkinson’s disease, Alzheimer’s disease and type II diabetes (Vafai and Mootha, 2012; Ploumi *et al*., 2017; Metaxakis *et al*., 2018). Mitophagy has also been shown to be important for developmental transitions in which the mitochondrial proteome needs to be drastically remodeled (Schweers *et al*., 2007; Sin *et al*., 2016; Esteban-Martínez *et al*., 2017). Known mitophagic mechanisms all involve a receptor protein which interacts with mitochondria on the one hand, and engages the autophagic machinery via LC3/ Atg8 proteins and ancillary factors in response to specific cues, on the other hand (Abeliovich and Dengjel, 2016). In mammalian cells, known mitophagy receptors include outer mitochondrial membrane (OMM) cardiolipin, NIX, BNIP3, NDP52, optineurin, prohibitin 2, TAX1BP1, FUNDC1 and Bcl2-L-13 (Schweers *et al*., 2007; Bellot *et al*., 2009; Novak *et al*., 2010; Liu *et al*., 2012; Lazarou *et al*., 2015; Murakawa *et al*., 2015; Wei *et al*., 2017), all of which function under different induction conditions and in different cell types. In budding yeast cells, a single OMM protein, Atg32, functions as the only known mitophagy receptor (Kanki *et al*., 2009; Okamoto *et al*., 2009). Current thinking envisions the selective activation of a mitophagy receptor on specific mitochondria, which leads to specific engulfment and degradation of those mitochondria. Thus, it is surprising that mitophagic processes, from yeast to man, exhibit protein level selectivity. We previously demonstrated, using SILAC-based proteomics and additional analyses, that different mitochondrial matrix proteins exhibit different mitophagic trafficking rates in yeast, and that this phenomenon correlates with physical segregation of the different proteins within the mitochondrial network (Abeliovich *et al*., 2013). Similar observations were reported in human cells (Hämäläinen *et al*., 2013; Burman *et al*., 2017) and in Drosophila (Vincow *et al*., 2013, 2019). More recently, we found that protein phosphorylation in the mitochondrial matrix plays a role in determining intra-mitochondrial mitophagic selectivity. Briefly, we discovered that mitophagic trafficking of the reporter protein Mdh1-GFP depends on its phosphorylation status, and that phosphorylation of the relevant residues requires the function of the mitochondrial matrix kinases Pkp1 and Pkp2, acting under the regulation of the Aup1/Ptc6 phosphatase, which had previously been implicated in mitophagy (Tal *et al*., 2007; Kolitsida *et al*., 2019).

Pkp1, Pkp2 and Aup1 have also been shown to regulate the phosphorylation of Pda1, the yeast E1α subunit of the mitochondrial pyruvate dehydrogenase complex (PDC) (Krause-Buchholz *et al*., 2006; Gey *et al*., 2008; Guo *et al*., 2017a). One simple explanation for the data would be that the pyruvate dehydrogenase reaction, which generates acetyl-coA from pyruvate in the mitochondrial matrix, is required for mitophagy or for mitophagic selectivity. However, several problems arise with this hypothesis. For example, in the regulation of pyruvate dehydrogenase activity, Pkp1 and Pkp2 function antagonistically relative to Aup1: they are required for phosphorylation of S313 on Pda1, which inactivates enzymatic activity, while Aup1 mediates dephosphorylation of this residue and activation of the enzyme (Guo *et al*., 2017a). In contrast, in the regulation of Mdh1-GFP mitophagic trafficking, Pkp2 and Pkp1 function together with Aup1: both the kinases and the phosphatase are required for efficient mitophagy of several reporters, with the kinases functioning downstream of Aup1, as Pkp2 overexpression suppresses the *aup1Δ* phenotype (Kolitsida *et al*., 2019).

To clarify and reconcile the relationship between the functions of Pkp1, Pkp2, and Aup1 in mitophagy versus their function in Pda1 regulation, we have now examined the potential role of the PDC in generating mitophagic selectivity. Surprisingly, the results of this investigation define a novel signaling cascade that is regulated by the PDC, but is uncoupled from its known enzymatic activity.

## Results

### Deletion of *pda1* has a marked effect on mitophagic selectivity, which is uncoupled from pyruvate dehydrogenase activity

Since Aup1, Pkp1 and Pkp2 have been identified as pyruvate dehydrogenase kinase phosphatases and kinases (Krause-Buchholz *et al*., 2006; Gey *et al*., 2008; González *et al*., 2013; Guo *et al*., 2017a), we wanted to understand the relationship between their role in regulating mitophagic selectivity, and their role in regulating pyruvate dehydrogenase activity. Surprisingly, we found that deletion of the *pda1* gene, which encodes the E1α subunit of the pyruvate dehydrogenase complex (PDC), has a major negative impact on the mitophagic trafficking of 5 different mitochondrial matrix proteins (Figure 1) when measured using the free GFP release assay. In contrast, the *pda1Δ* mutation had no effect on the mitophagic trafficking of an artificial mitophagy reporter, mtDHFR-GFP, as we previously reported for the *pkp1Δ pkp2Δ* double mutant (Kolitsida *et al*., 2019). This result implies that the *pda1Δ* mutation does not block mitophagy per se, but rather affects the sorting of individual proteins to a mitophagy-targeted mitochondrial sub-population, based on native protein-protein interactions that are not accessible to the artificial DHFR-based reporter. In addition, deletion of *pda1* had no effect on the levels of Pkp1, Pkp2, or Aup1 (Supplementary Figure 1).

**Figure 1.**
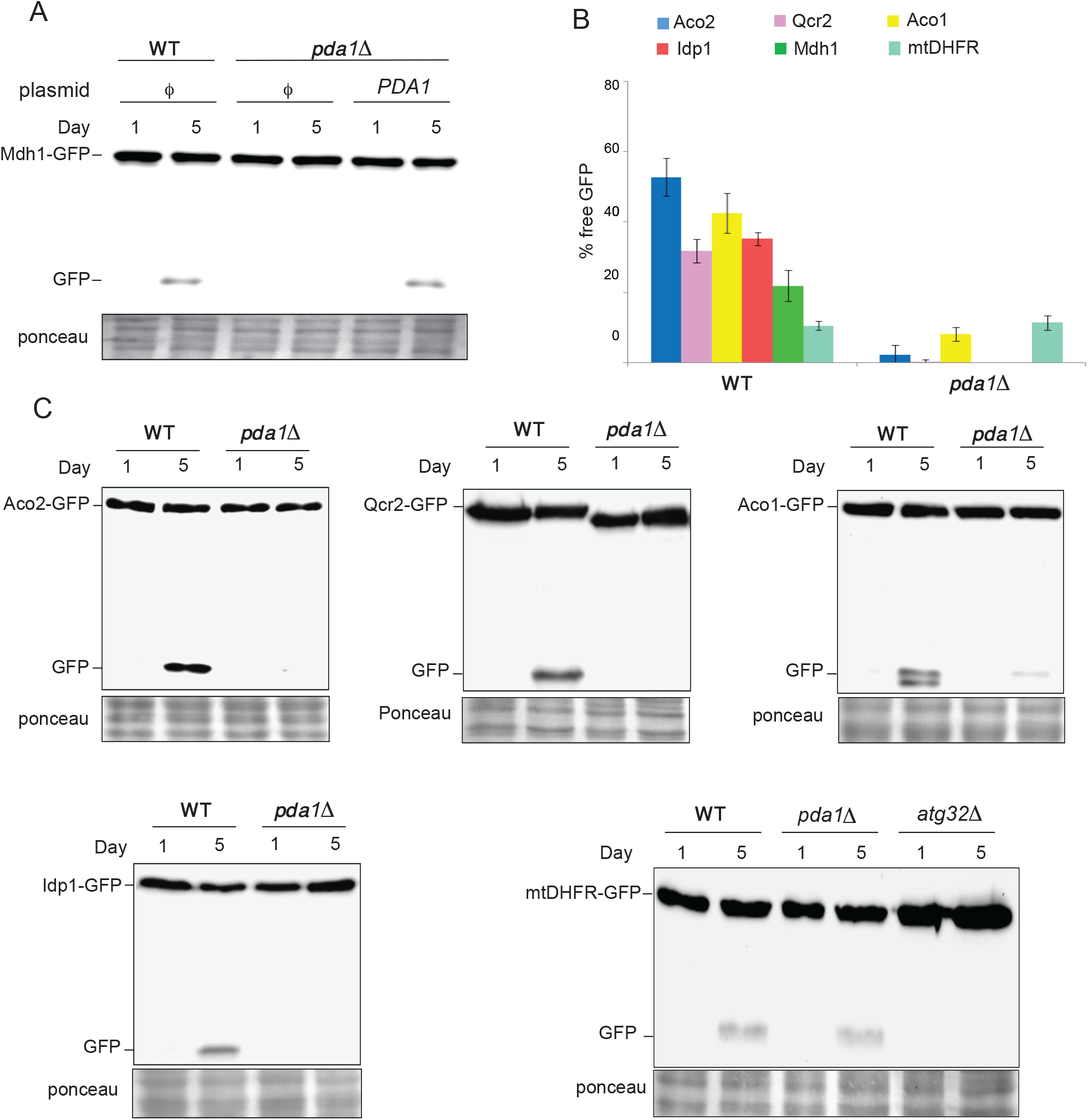
Loss of *pda1* affects mitophagic trafficking and selectivity of mitochondrial matrix proteins. WT (HAY75) and *pda1Δ* (PKY832) cells expressing the indicated GFP-tagged mitochondrial matrix proteins were grown in SL medium as indicated in “Methods”. Samples were taken on days 1 and 5, and protein extracts were analyzed by anti-GFP immunoblotting to determine the percent of signal converted to free GFP. **A**. Loss of *pda1* strongly attenuates mitophagic trafficking of Mdh1-GFP and this can be complemented by expressing pda1 from a plasmid. **B**. Loss of *pda1* strongly attenuates the mitophagic trafficking of endogenous GFP chimeras of five different mitochondrial matrix proteins, but not of the ‘artificial’ mtDHFR-GFP reporter. **C**. Representative immunoblots for the data summarized in (B). Data are shown from 3 independent biological replicates +/- standard deviation. *φ* denotes empty vector.

The most straightforward explanation for this result would be that the pyruvate dehydrogenase reaction itself, i.e. the formation of acetyl-CoA from pyruvate with concomitant reduction of NAD^+^, is required for efficient mitophagy. However, deletion of the mitochondrial pyruvate carrier subunit *MPC1* had no effect on mitophagic trafficking (Figure 2A, B), even though this mutation prevents entry of pyruvate into mitochondria (Herzig *et al*., 2012). We also tested whether the role of Aup1 in mitophagy is limited to the dephosphorylation of pda1 at position Ser313 (the residue regulated by Pkp1/2 and Aup1; (Gey *et al*., 2008; Guo *et al*., 2017a)). To this end, we determined whether deletion of *AUP1* can affect mitophagic trafficking in cells where the sole copy of *pda1* encodes a serine to alanine mutation at position 313. As shown in Figure 2C and 2D, the pda1^S313A^ mutant can weakly complement deletion of the WT protein for mitophagic trafficking of the Mdh1-GFP reporter. However, this residual mitophagic trafficking is completely lost in cells which also lack Aup1, indicating that Aup1 regulates mitophagic trafficking of Mdh1 independently of the phosphorylation state of serine 313 on pda1. Finally, we tested whether catalytically inactive point mutations in the E1 and E2 subunits affect the mitophagic trafficking of the Mdh1-GFP reporter. For the E1α subunit pda1, we chose the R322C mutation, which was previously shown to completely abrogate catalytic activity (Drakulic *et al*., 2018). As shown in Figure 2E and 2F, although this point mutation caused a significant decrease in mitophagic trafficking of Mdh1-GFP, it still allowed a reproducible signal of free GFP under mitophagy inducing conditions, which was significantly higher than that observed in the *pda1Δ* deletion, and does not phenocopy the deletion. More importantly, when we introduced the un-lipoylatable K75R mutation in the E2 subunit Lat1 (Lawson *et al*., 1991; Gey *et al*., 2014), expression of this catalytically inactive mutant subunit led to a clear increase in mitophagic trafficking of Mdh1-GFP-completely opposite from the result expected if the *lat1Δ* phenotype was the result of a block in PDC catalytic activity (Fig. 2G, H). Neither *lat1Δ* nor *lat1*^K75R^ had any effect on the mitophagic trafficking of mtDHFR-GFP (Supplementary Figure 2). Collectively, these results indicate that the effects of the *pda1Δ* and *lat1Δ* mutations on mitophagic trafficking cannot simply be explained by the loss of PDC enzymatic activity, and are uncoupled from the phosphorylation state of Serine 313 on pda1.

**Figure 2.**
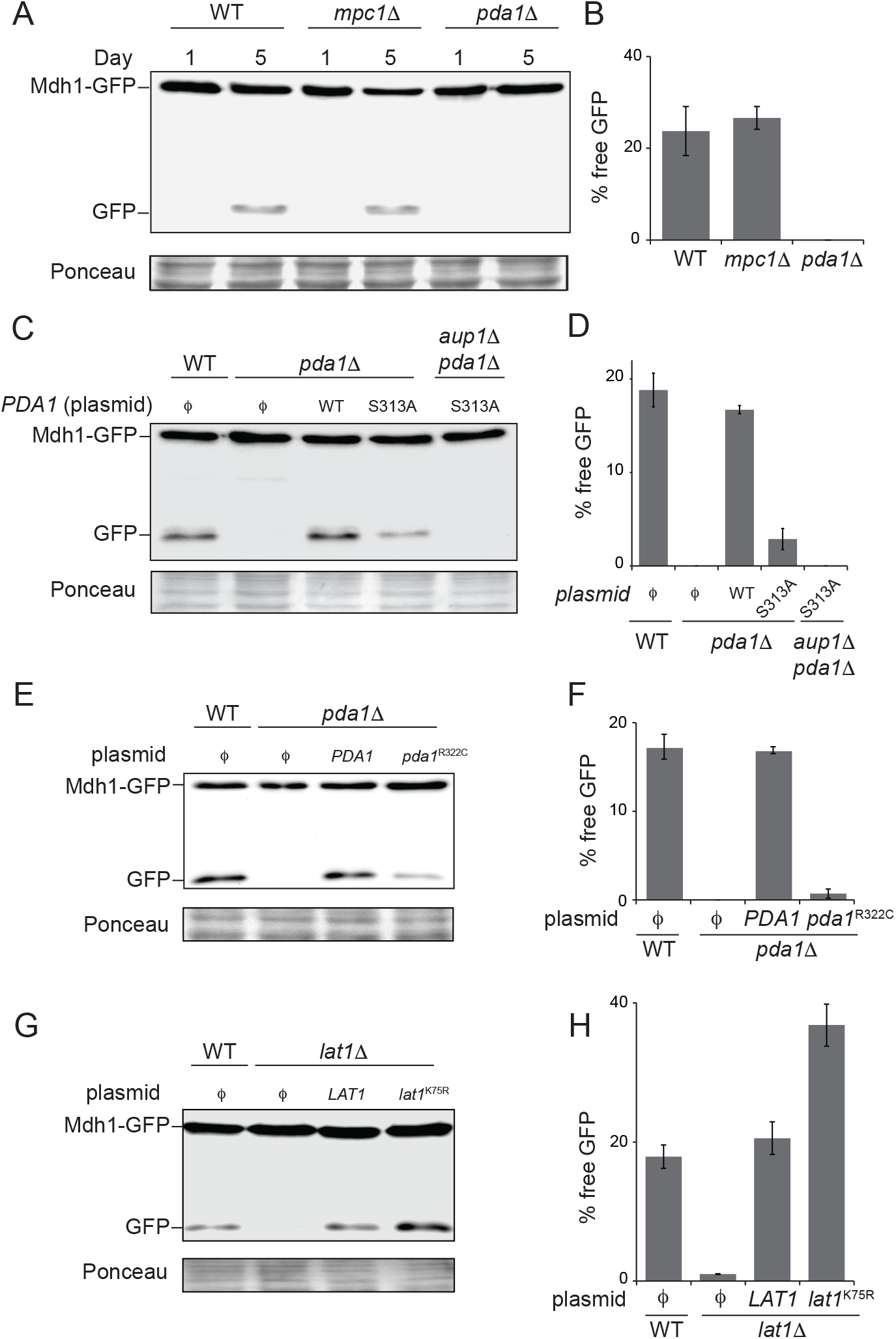
The effects of the *pda1* deletion on mitophagy are not mediated through the pyruvate dehydrogenase reaction, and mitophagic trafficking can be modulated independent of S313 phosphorylation. **A**. Loss of *MPC1* has no effect on mitophagic trafficking of Mdh1-GFP. WT (PKY974), *mpc1Δ* (PKY2104) and *pda1Δ* (PKY2023) cells expressing Mdh1-GFP were grown in SL medium for the indicated amounts of time and protein extracts were generated. The extracts were analyzed by immunoblotting with α-GFP antibodies. **B**. Quantification of the data in A. **C**. Cells expressing the S313A pda1 point mutant still exhibit an *aup1*-dependent mitophagic trafficking phenotype despite the absence of phosphoserine 313. PKY2023 *(pda1Δ)* and PKY2204 (*aup1Δ pda1Δ*) Cells were grown in SL and handling was as in (A), but only the day 5 samples are shown. **D**. Quantification of the data in C. **E**. The catalytically inactive *pda1*^R322C^ mutant can support partial mitophagic trafficking of Mdh1-GFP. Only day 5 samples are shown for brevity. **F**. Quantification of the data in (E). **G**. The unlipoylatable *lat1*^K75R^ mutant shows increased mitophagic trafficking of Mdh1-GFP, relative to WT. Only day 5 samples are shown, for brevity. **H**. Quantification of the data in F. All quantifications are presented as averages of 3 independent biological replicates with standard deviation depicted as error bars. *φ* denotes empty vector.

### The PDC affects phosphorylation of Mdh1-GFP via regulation of Pkp1 and Pkp2

The extent of the decrease in mitophagic trafficking of mitochondrial matrix GFP chimeras, which we observe in *pda1Δ* cells (Figure 1A), is very similar to the phenotype which we had previously reported for the *pkp1Δ pkp2Δ* double mutant (Kolitsida *et al*., 2019). Those previous studies showed that Mdh1-GFP is strongly hypo-phosphorylated in *pkp1Δ pkp2Δ* double mutants (Kolitsida *et al*., 2019). An alternative hypothesis that we therefore wanted to explore, was that the PDC reciprocally regulates its associated kinases and phosphatases, by virtue of their physical association (Krause-Buchholz *et al*., 2006; Breitkreutz *et al*., 2010; Guo *et al*., 2017a), as such interactions with the PDC could potentially regulate the enzymatic activities of Aup1, Pkp1 and Pkp2. If the effect of the *pda1Δ* mutation reflects a reciprocal regulation of Aup1, Pkp1 and Pkp2 by the PDC, we would expect that loss of pda1 would similarly affect the phosphorylation of Mdh1-GFP. Indeed, we find that Mdh1-GFP phosphorylation is strongly attenuated in *pda1Δ* cells, and that this phenotype can be complemented by expressing pda1 from a plasmid (Figure 3A, B). Similarly, knockout of *pda1* also led to hypo-phosphorylation of Qcr2-GFP and Aco1-GFP under these conditions (supplementary figures 3 and 4). It was previously shown that Aup1 and Pkp1 stably interact with the PDC (Krause-Buchholz *et al*., 2006; Breitkreutz *et al*., 2010; Guo *et al*., 2017a). To determine whether the PDC might function upstream of the kinases in regulating mitophagic trafficking of Mdh1-GFP, we tested if overexpression of Pkp1, Pkp2 or both kinases together, might rescue the mitophagy defect observed in *pda1Δ* cells. As shown in Figure 3C and 3D, co-overexpression of Pkp1 together with Pkp2, but not each kinase separately, caused a near-complete rescue of the *pda1Δ* mitophagy phenotype. Importantly, deletion of *pda1* did not affect the levels of Pkp1, Pkp2, or Aup1 (supplementary Figure 1), indicating that loss of pda1 does not destabilize these proteins or otherwise affect their expression. In addition, we were able to show a similar suppression pattern for overexpression of Pkp1 and Pkp2 in *aup1Δ* cells, supporting the hypothesis that the PDC regulates Pkp1 and 2 via an effect on Aup1 (Figure 3E, We previously showed that the mitophagy phenotype of the *pkp2Δ* mutation could be suppressed in cells expressing the Mdh1^T199D^-GFP reporter, which mimics phosphorylation of Mdh1 at residue 199 (Kolitsida *et al*., 2019). If the effects of the *pda1Δ* mutation are due to an effect of the PDC on Pkp1 and Pkp2, then we expect that, similarly, the Mdh1^T199D^-GFP reporter would show at least a partial suppression in the *pda1Δ* background. As shown in Figure 3G and 3H, expression of the Mdh1^T199D^-GFP variant, but not the Mdh1^T199A^-GFP mutant, efficiently bypassed the Mdh1 mitophagic trafficking defect block in the *pda1Δ* mutant. Taken together, these results strongly support the argument that pda1 affects mitophagy by regulating the phosphorylation state of the reporter.

**Figure 3.**
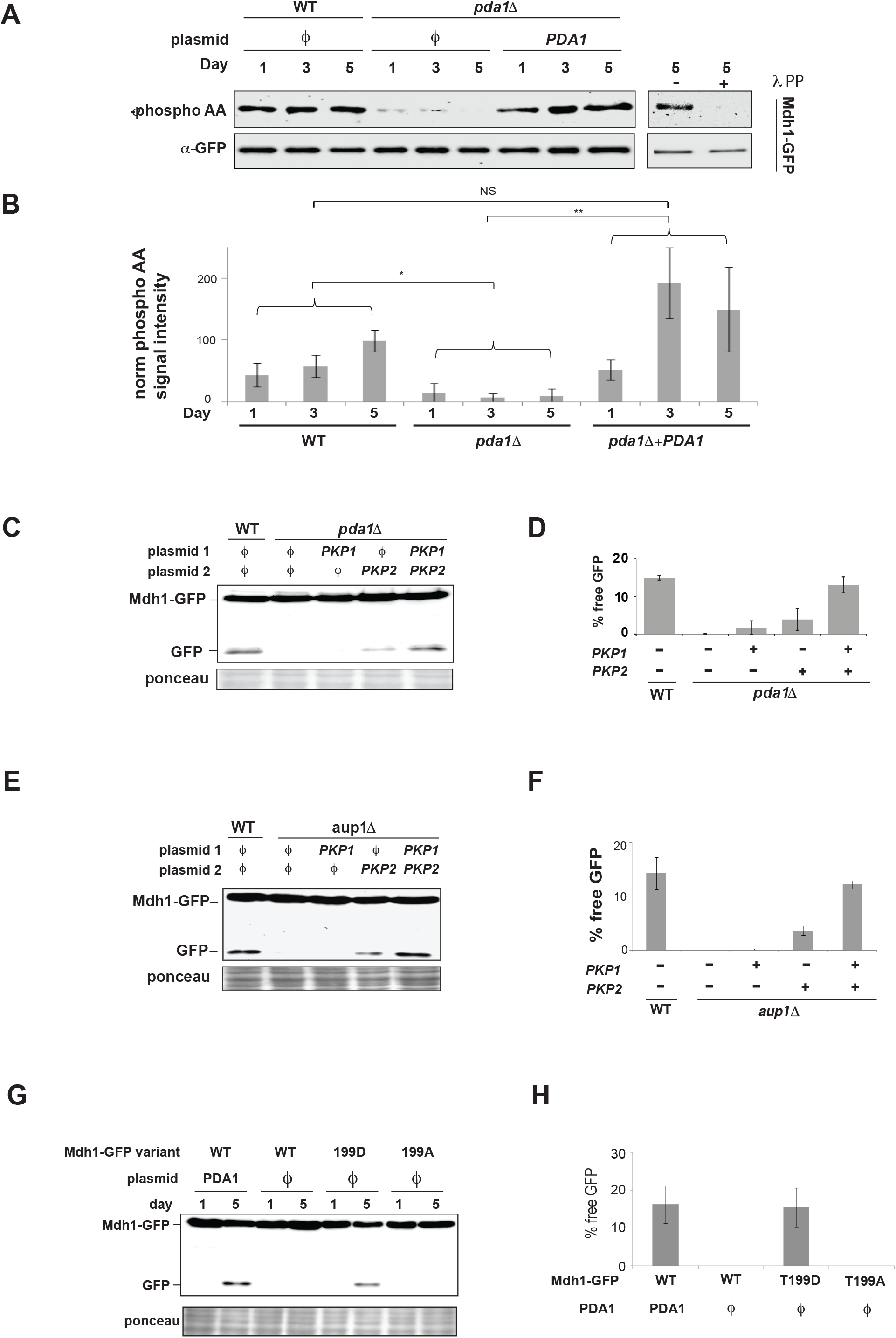
The PDC regulates phosphorylation of Mdh1. **A**. The *pda1Δ* mutation causes hypophosphorylation of Mdh1-GFP. WT (PKY1089) and *pda1Δ* (PKY1093) cells expressing Mdh1-GFP were grown in SL medium for the indicated amounts of time and protein extracts were generated and immunoprecipitated with α-GFP antibody. The immunoprecipitates were then probed with α-phosphoamino acid antibody (top panel) and α-GFP antibody (bottom panel). Right panel: day 5 samples were treated overnight with lambda phosphatase or mock treated as described in “Methods”, and analyzed by immunoblotting. **B**. Quantification of the data in (**A**). The results of 3 independent biological repeats were quantified by normalizing the α-phosphoamino acid signal to the GFP signal, and averaging. **C**. Co-overexpression of pkp1 and pkp2 completely suppresses the mitophagy defect of *pda1Δ* cells. WT (PKY974) and *pda1Δ* (PKY2023) cells expressing Mdh1-GFP as well as the indicated plasmids, were grown in SL medium for 5 days and protein extracts were generated on day 5. The extracts were analyzed by immunoblotting with *α*-GFP antibodies. **D**. Quantification of the data in (C). The results of 3 independent biological repeats were quantified by calculating the % free GFP relative to the total signal (N=3) and averaging. **E**. Co-overexpression of pkp1 and pkp2 completely suppresses the mitophagy defect of *aup1Δ* cells. WT (PKY974) and *aup1Δ* (HAY809) cells expressing Mdh1-GFP as well as the indicated plasmids, were grown in SL medium for 5 days and protein extracts were generated on day 5. The extracts were analyzed by immunoblotting with α-GFP antibodies. **F**. Quantification of the data in (E). The results of 3 independent biological repeats were quantified by calculating the % free GFP relative to the total signal (N=3) and averaging. **G**. Expression of Mdh1^T199D^-GFP, but not Mdh1^T199A^-GFP bypasses the effect of the *pda1Δ* mutation. PKY2357 cells expressing Mdh1-GFP, Mdh1^T199D^-GFP, and Mdh1^T199A^-GFP were grown as described above. Protein extracts were generated on days 1 and 5 and analyzed as above. **H**. Quantification of the data in (G). All experiments represent an average of at least 3 independent biological repeats and statistical analysis was carried out as described in Materials and Methods. NS= not significant; Asterisks denote significant (p< 0.05) differences.

Furthermore, co-overexpression of Pkp1 and Pkp2 increases the phosphorylation of Mdh1-GFP in *pda1Δ* cells, as would be expected if the PDC was modulating Pkp1/2 activity towards downstream targets (Figure 4A, 4B), again supporting our finding that it is not PDC catalytic activity which is important for the mitophagic trafficking of Mdh1-GFP.

**Figure 4.**
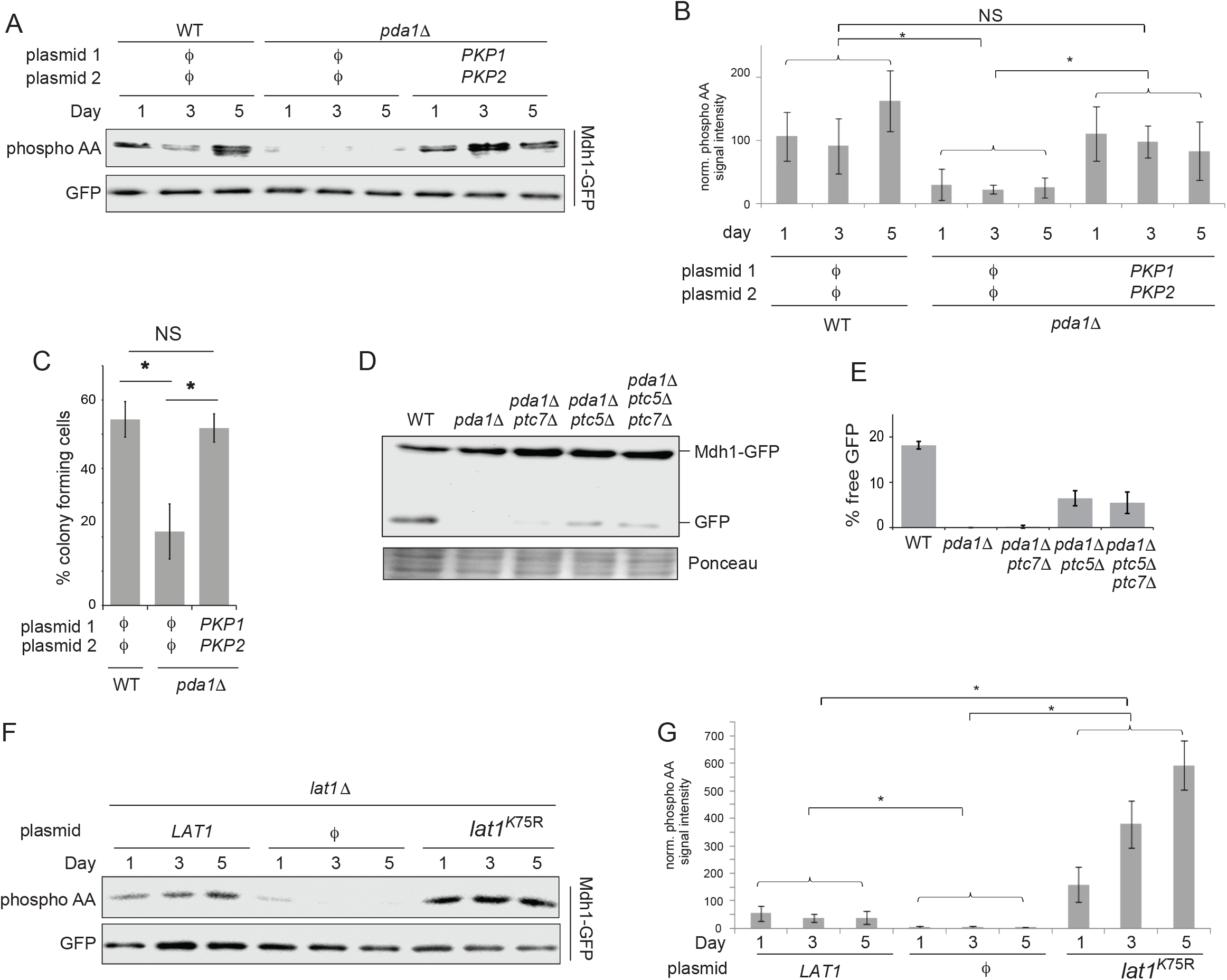
Mdh1-GFP phosphorylation and mitophagic trafficking are regulated by the PDC complex independent of the pyruvate dehydrogenase reaction. **A**. Co-overexpression of pkp1 and pkp2 suppresses the Mdh1-GFP phosphorylation defect in *pda1Δ* cells. WT (PKY1089) and *pda1Δ* (PKY1093) cells expressing Mdh1-GFP as well as the indicated plasmids were grown in SL medium for the indicated amounts of time and protein extracts were generated and immunoprecipitated with α-GFP antibody. Phosphorylation of Mdh1-GFP was detected using α-phosphoamino acid antibody and total GFP signal in the immunoprecipitate was observed by immunoblotting with α-GFP antibody. *φ* denotes empty vector. **B**. Quantification of the data in (A). The results of 3 independent biological repeats were quantified by normalizing the *α*-phosphoamino acid signal to the GFP signal, and averaged. **C**. pkp1 and pkp2 have a physiological function independent of pda1 phosphorylation. Cells (HAY75 and PKY832) were grown to late log phase in SD medium, washed and equal cell numbers were resuspended in SL medium. Cell viability was measured by plating diluted aliquots on rich YPD medium after 5 days of incubation in SL medium. Clearly, cooverexpression of pkp1 and pkp2 completely suppresses the defect in *pda1Δ* cells (N=3, error bars denote standard deviation), * denotes p<0.05 (t-test). NS - not significant. **D**. Deletion of PTC5 partially suppresses the *pda1Δ* mitophagy defect. Mdh1-GFP-expressing WT (PKY974), *pda1Δ (PKY2023)*, and *pda1Δ* cells harboring co deletions of *PTC7* (PKY2234), *PTC5* (PKY2239) or both (PKY2286), were incubated in SL medium for 5 days and protein extracts were generated. The extracts were analyzed by immunoblotting with *α*-GFP antibodies. **E**. Quantification of 3 independent biological repeats of the experiment shown in (D). **F**. The catalytically inactive Lat1^K75R^ mutant causes massive hyperphosphorylation of Mdh1-GFP. Mutant *lat1Δ* cells (PKY2298) expressing the indicated plasmids as well as Mdh1-GFP, were grown in SL medium for the indicated amounts of time and protein extracts were generated and immunoprecipitated with α-GFP antibody. Phosphorylation of Mdh1-GFP was detected and quantified as in (A, B). **G**. Quantification of the data in (F). All experiments represent an average of at least 3 independent biological repeats and statistical analysis was carried out as described in Materials and Methods. NS= not significant; Asterisks denote significant (p< 0.05) differences.

We were also able to demonstrate a clear physiological function for Pkp1 and Pkp2, that is uncoupled from the phosphorylation of the E1α subunit: While *pda1Δ* cells show a marked reduction in the percentage of colony forming cells following a prolonged incubation in respiratory medium, co-overexpression of Pkp1 and Pkp2 fully suppressed this phenotype (Figure 4C and Supplementary Figure 5). This result demonstrates that the two kinases have a physiological function other than the regulation of PDC catalytic activity, and at the same time the epistasis relationships indicate-like our other results-that the PDC seems to act as an upstream regulatory element in the(se) novel signaling pathway(s), independent of its catalytic activity. If pda1 regulates Mdh1-GFP phosphorylation via regulation of the activities of Pkp1 and Pkp2, then co-deletion of the opposing phosphatase should partially reverse this defect, as any residual phosphorylation activity would now be amplified by the loss of the opposing phosphatase activity. Our previous work demonstrated that Aup1/Ptc6 cannot be opposing Pkp1/Pkp2 in regulating mitophagic trafficking, since 1) deletion of *AUP1* leads to a phenotype that is similar to the phenotypes of *PKP1Δ* and *pkp2Δ* on mitophagic trafficking, and 2) aup1/ Ptc6 functions upstream of pkp1 and pkp2 in regulating Mdh1-GFP phosphorylation and mitophagic trafficking (Kolitsida *et al*., 2019). We therefore tested the effects of deleting the mitochondrial phosphatases *PTC5* and *PTC7* (Guo *et al*., 2017a, 2017b) on Mdh1-GFP mitophagic trafficking, in the absence of pda1. As shown in Figure 4D and 4E, deletion of *PTC5*, and to a much lesser extent deletion of *PTC7*, both led to some generation of free GFP in *pda1Δ* cells, while the phenotype of the combined *ptc5Δ pdc7Δ* double deletion seems identical to the *ptc5Δ* phenotype. This result further supports the hypothesis that the PDC regulates the phosphorylation state of Mdh1-GFP, and suggests that Ptc5 is the main mitochondrial phosphatase which antagonizes the function of pkp1 and pkp2 in regulating mitophagic trafficking of Mdh1-GFP (see discussion).

To further test the possibility that conformational changes in the PDC may regulate pkp1 and pkp2 activity, we asked whether the catalytically inactive *lat1*^K75R^ mutation, which increases mitophagic efficiency of Mdh1-GFP (Figure 2G, H), also increased phosphorylation of this reporter. As shown in Figures 4F and 4G, deletion of *LAT1* decreased phosphorylation of Mdh1-GFP, while the K75R point mutation drastically increased phosphorylation of the protein by 4-10 fold under these conditions, relative to the phosphorylation level of the reporter in WT control cells. Again, this result supports a conformation-dependent regulation of pkp1/2 by the PDC, which is decoupled from PDH catalytic activity.

### Deletion of *pda1* affects the distribution of Mdh1 in the mitochondrial network, relative to an artificial mtRFP reporter

We previously showed that deletion of aup1, which functions upstream of pkp2 in the cascade regulating mitophagic selectivity, leads to increased segregation of Mdh1-GFP relative to mtRFP (which does not make native protein protein interactions with matrix proteins, and is therefore not expected to be segregated in the network), within the mitochondrial matrix (Kolitsida *et al*., 2019). This was interpreted as indicating that native protein protein interactions between matrix proteins, which are not accessible to the mtRFP reporter, are important in segregating endogenous mitochondrial matrix GFP chimeras into mitophagy-bound or mitophagy-blocked mitochondrial populations. To test whether the deletion of *pda1* similarly affects mitochondrial matrix heterogeneity, we compared the signal overlap between mtRFP and Mdh1-GFP in wild type and *pda1Δ* cells, over the first two days of incubation in SL medium (Figure 5). Wild-type cells show a slight, statistically insignificant decrease in signal overlap between day 1 and day 2 of the incubation. The *pda1Δ* mutant however, showed a statistically significant, ~40% reduction in signal overlap of Mdh1-GFP with mtRFP on day 2 relative to day 1 (p<0.0001), similar to what we previously observed for *aup1Δ* cells (Kolitsida *et al*., 2019). Importantly, co-overexpression of pkp1 and pkp2 in the *pda1Δ* cells suppressed the decrease in signal overlap between days 1 and 2 of the incubation. In addition, we observed a mild mitochondrial morphology phenotype in the *pda1Δ* strain, which appeared on day 2, in which both the red and the green signals appeared to be more clumped into a smaller number of larger foci relative to both WT and kinase-overexpressing strains.

**Figure 5.**
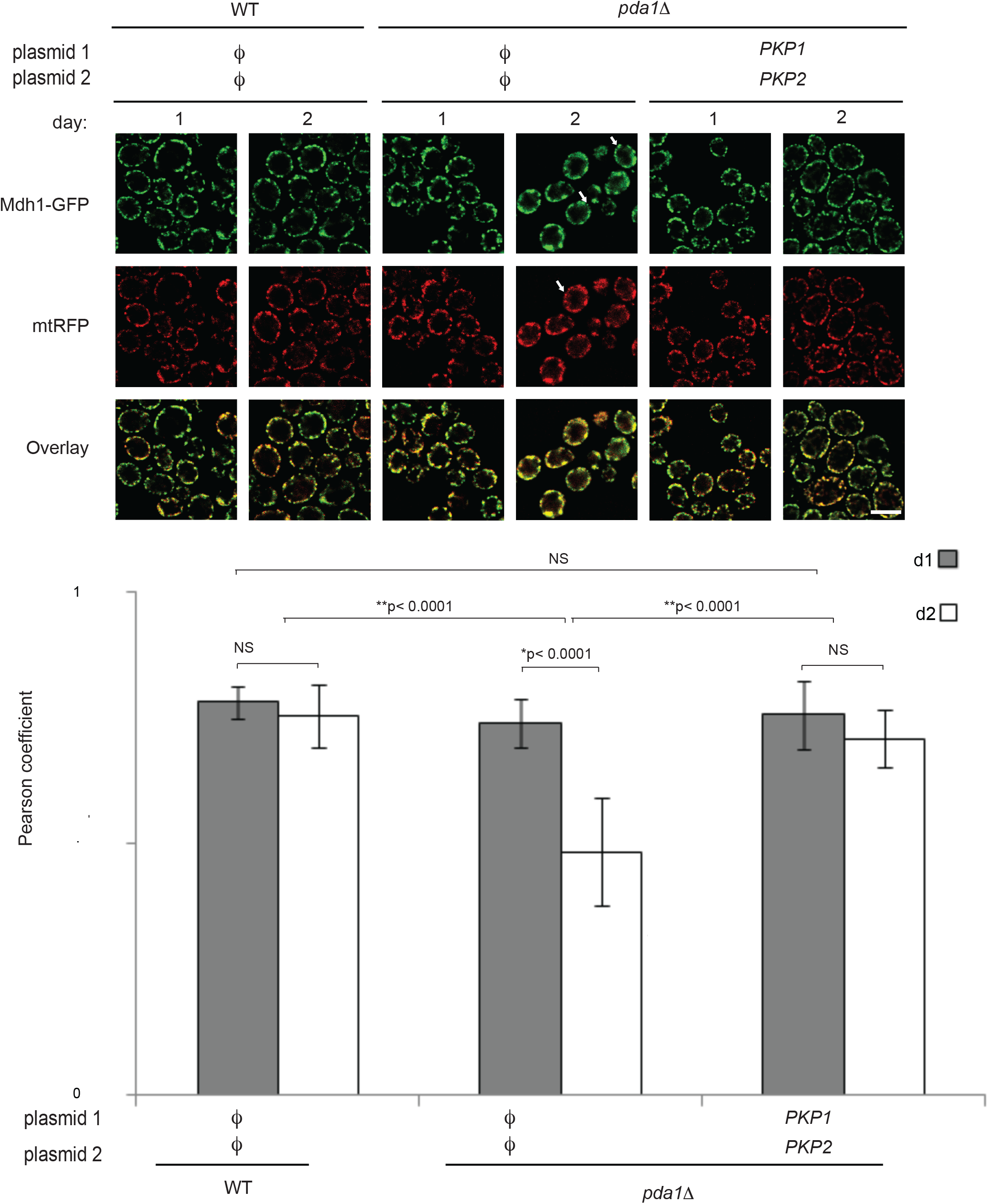
Deletion of *pda1* affects the intra-mitochondrial distribution of Mdh1-GFP. **A.** Deletion of *pda1* affects distribution of Mdh1-GFP within the mitochondrial matrix. Cells (PKY974 and PKY2023) co-expressing Mdh1-GFP from the endogenous *MDH1* promoter and mtRFP (which does not make native protein-protein interactions with matrix proteins, and is therefore not expected to be segregated within the network) were imaged on days 1 and 2 of the incubation (to avoid interference from vacuolar signal from day 3 and up) and the overlap between the red and green channels was quantified as described in ‘Materials and Methods’. White arrows indicate individual examples of differential distribution of fluorescence between the two channels. **B**. Graph showing quantification of the overlap between the two channels in WT and in *pda1Δ* cells, and indicating statistical significance (NS, not significant). Results are representative of three biological replicates. Scale bar =5 μm.

### The Lat1^K75R^ mutation affects physical interactions between the PDC, aup1 and pkp1/2

If The PDC affects mitophagy through its physical interactions with aup1, pkp1 and pkp2, we expect that the Lat1^K75R^ mutant, which has a clear effect on mitophagy, would affect the known physical interactions between aup1, pkp1, pkp2 and the PDC. As shown in Figure 6, we were able to recapitulate the interaction between aup1 and the PDC under our experimental conditions. We observe that the interaction between aup1 and pda1 is weak on day 1 of our experimental timeline, but becomes much stronger on day 3. At the same time, we observe a shift in the mobility of aup1-HA to slightly faster migration in the day 3 samples, relative to day 1. Strikingly, the Lat1^K75R^ mutation abrogates any observable co-immunoprecipitation between aup1 and pda1, at both time points. This demonstrates that the conformational change induced by the K75R mutation affects the aup1-PDC interaction, indicating a potential mechanistic explanation for the up-regulation of Mdh1-GFP mitophagic trafficking and phosphorylation in cells harboring this mutation.

**Figure 6.**
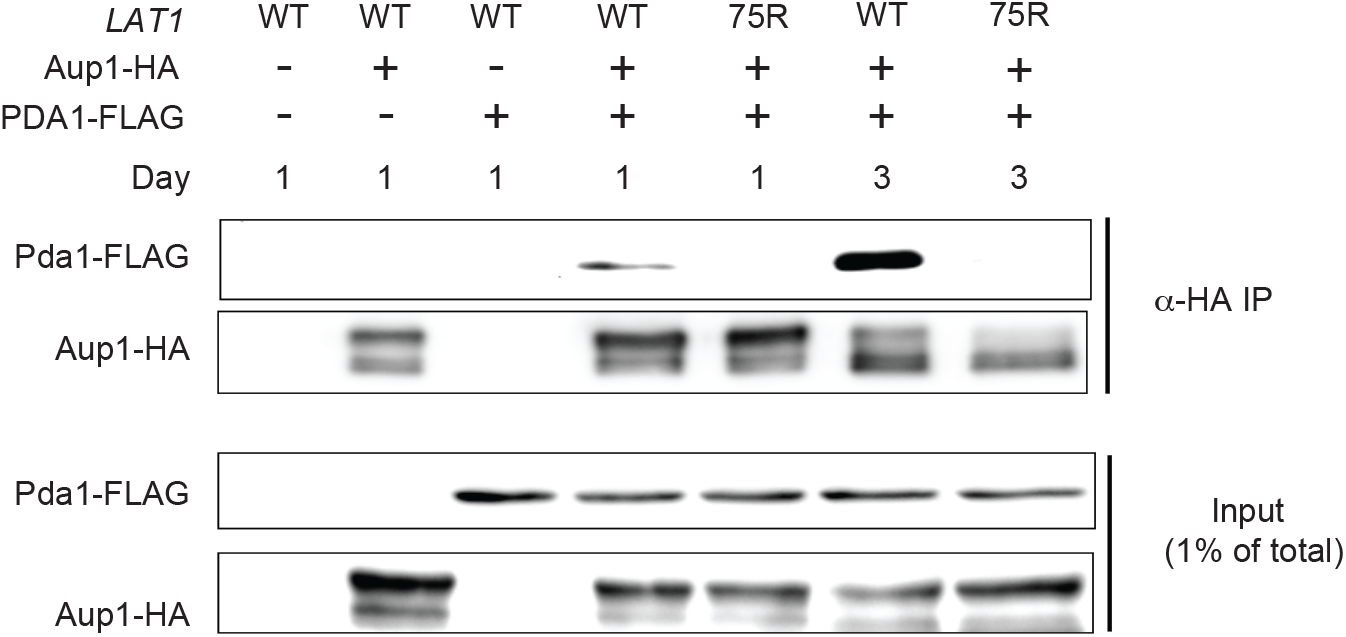
The Lat1^K75R^ mutation affects the physical interaction between aup1 and the PDC. Cells deleted for *LAT1* (PKY2498) and expressing aup1-HA and pda1-FLAG as well as the indicated *LAT1* variant (empty vector, WT or K75R) were incubated in SL medium for 1 or 3 days as indicated and samples were analyzed by immunoprecipitation with α-HA antibodies and immunoblotted for HA or FLAG epitopes as indicated.

### The Lat1^K75R^ mutation affects protein phosphorylation in the mitochondrial matrix

Our experiments demonstrate a previously undescribed regulation of mitochondrial protein phosphorylation by the PDC, via pkp1 and pkp2. Since the Lat1^K75R^ mutation appears to increase phosphorylation of the Mdh1-GFP reporter, we wanted to determine whether this is a more global effect on protein phosphorylation in the matrix. To this end, WT versus Lat1^K75R^ expressing cells were differentially labelled with heavy and light isotope-substituted arginine and lysine. The cultures were collected on day 3 of an incubation in synthetic lactate medium (SL), the harvested cells were mixed in equal amounts, and mitochondria were enriched by differential centrifugation. As shown in Figure 7 and in Supplementary Table IV, we were able to identify at least 21 phosphorylation sites from 19 different mitochondrial matrix proteins, whose phosphorylation state was significantly and reproducibly affected by the deletion of *pda1*. These results demonstrate that PDC-mediated regulation of protein phosphorylation in the mitochondrial matrix is not limited to Mdh1. However, the distribution of the results also clearly indicates that the mutation causes both hypo-phosphorylation as well as hyperphosphorylation, depending on the phosphorylation site and the specific protein. We therefore suggest that the effect of the PDC on pkp1 and pkp2 - dependent phosphorylation is not strictly on/off, but rather an effect on kinase selectivity. Interestingly, the proteins which showed significant hyper-phosphorylation in the Lat1^K75R^ mutant included the mitophagy receptor Atg32. We also identified significant hits in previously undocumented phosphorylation sites in PDC subunits.

**Figure 7.**
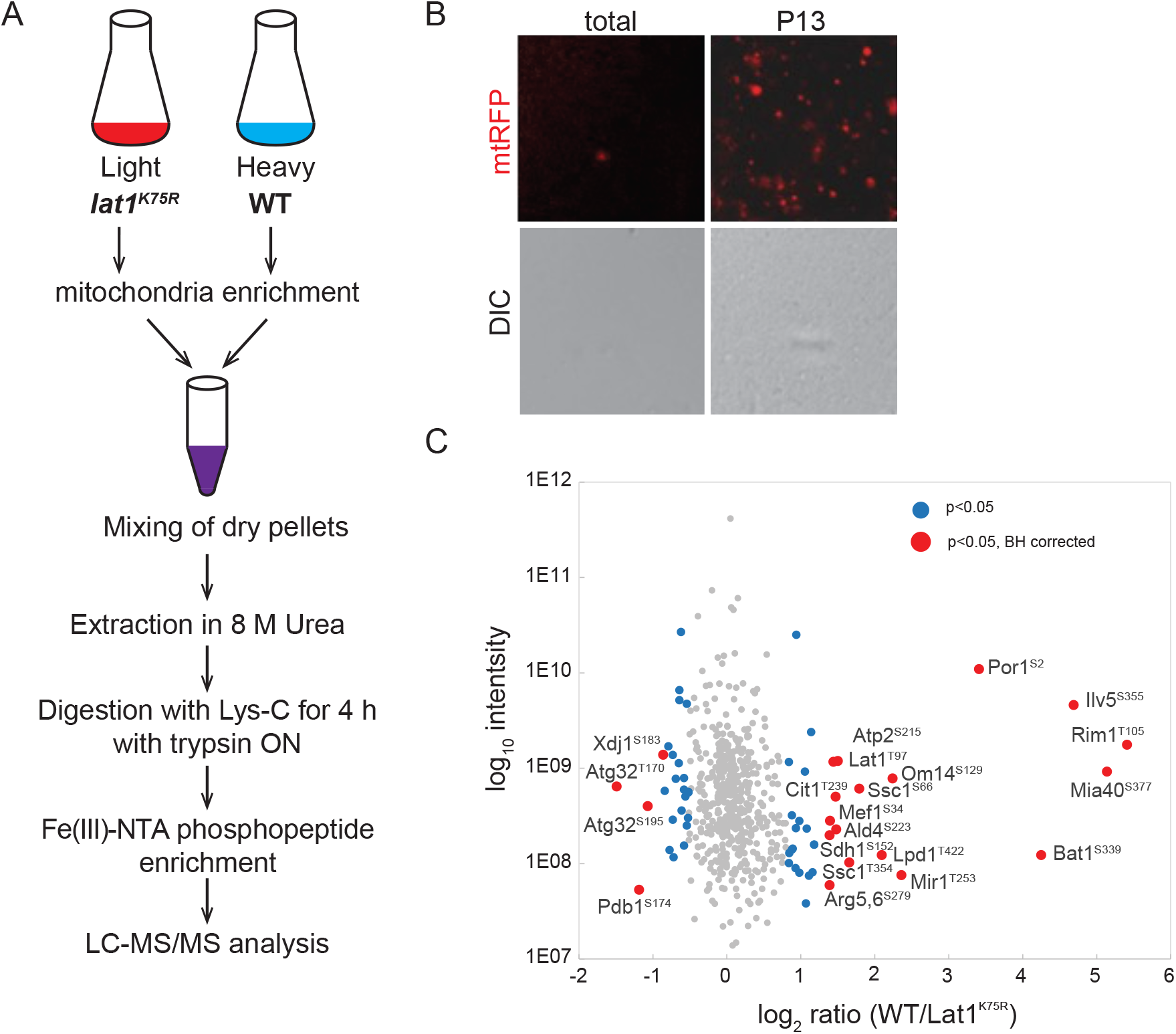
The *lat1*^K75R^ mutation has diverse effects on protein phosphorylation in the mitochondrial matrix. **A**. Overview of the methodology. WT (PKY2154) cells and *pda1Δ* (PKY2196) cells expressing mtRFP were grown for 2 days in SL medium containing heavy lysine+arginine (WT) or regular lysine+arginine (*pda1Δ*). Equivalent amounts of WT and *pda1Δ* cells were combined (see Materials and Methods) and cell extracts were generated by gentle cryomilling. **B**. The extract was further centrifuged at 13000 xg and the pellet was analyzed by microscopic examination (100x objective) to verify the enrichment of intact mitochondria relative to total extract. DIC images are included to verify the absence of intact cells. **C**. Quantitative SILAC MS/MS comparison of mitochondrial phosphopeptides identified in WT and *pda1Δ* mutants. Significant regulated sites are marked by color, p<0.05 in blue, p<0.05, BH-corrected in red. Significant hits after correction are annotated by protein. Phosphorylated positions are denoted in superscript.

## Discussion

In this study, we identified novel regulatory interactions between the pyruvate dehydrogenase complex and its attendant kinases. While previous studies, from the 1960’s to the present, have focused on the fact that PDKs regulate pyruvate dehydrogenase activity (Linn *et al*., 1969; Gey *et al*., 2008; Guo *et al*., 2017a; Cardoso *et al*., 2020), our data now demonstrate that the PDKs have additional substrates, and that their activity towards these non-PDC substrates can be reciprocally regulated by the PDC itself. This regulation may be mediated via allosteric interactions, as it was previously shown that pkp1 and pkp2 function as a heterodimer (Gey *et al*., 2008; Breitkreutz *et al*., 2010; Walter *et al*., 2022) and both aup1 and pkp1 stably interact with the PDC (Krause-Buchholz *et al*., 2006; Guo *et al*., 2017a). Hundreds of mitochondrial matrix proteins have been documented to undergo phosphorylation (Reinders *et al*., 2007; Kruse and Højlund, 2017; Renvoisé *et al*., 2017). However, the nature of the signaling cascades involved and their regulation has remained obscure. Our discovery that pkp1 and pkp2 can function to phosphorylate mitochondrial matrix proteins other than pda1 opens up the possibility that at least some of the documented phosphorylation events in the mitochondrial matrix are due to this novel regulatory axis. In support of this, we found by MS analysis that at least 19 additional mitochondrial matrix proteins show differential phosphorylation in response to the *lat1*^K75R^ mutation, under our working conditions (Figure 7). While this list is not comprehensive (many peptides were not identified in all three biological replicates, including the Mdh1 phosphopeptides; See supplementary table IV) it provides an indication for a more general effect of the PDC on mitochondrial phosphorylation. We previously showed a functional link between aup1 and the RTG retrograde signaling pathway (Journo *et al*., 2009) which transduces stress signals from the mitochondrial matrix to the nucleus (Liu and Butow, 2006), and this may account for the effects on non-matrix proteins such as Atg32. We hypothesize that some of this regulation of mitochondrial protein phosphorylation by the PDC can be ascribed to an allosteric effect of the PDC on aup1/Ptc6 and pkp1/2. As aup1 has been shown to interact with the PDC (Guo *et al*., 2017a), its phosphatase activity towards other substrates could be influenced by this stable physical interaction. In agreement with this, we can observe the PDC-aup1 interaction under our conditions, and we further demonstrate that it is strongly affected by the *lat1*^K75R^ mutation (Figure 6). Interestingly, the effects of the *lat1*^K75R^ mutant on matrix protein phosphorylation are diverse, with some sites being hypophosphorylated and others hyperphosphorylated in cells expressing the mutant, relative to WT cells (Figure 7). This may indicate that the regulation is not a simple on/off switch but rather a change in substrate specificity, suggesting that the association of pkp1/2 and aup1 with the PDC regulates substrate specificity, and not simply the activity of the kinases. In support of this, full suppression of the *pda1Δ* phenotype requires overexpression of both pkp1 and pkp2 (Figure 3C, D), suggesting that multiple phosphorylation (and potentially also de-phosphorylation) events are required, in effect altering the physical properties of matrix proteins and their mutual interactions. In addition, we find that the *lat1*^K75R^ mutation causes significantly increased phosphorylation of Atg32 (Figure 7 and Supplementary Table IV). In agreement our observation that the *lat1*^K75R^ mutant displays increased phosphorylation and mitophagic trafficking of Mdh1-GFP (Figure 2G and Figure 4F), phosphorylation of Atg32 was shown by others to regulate its mitophagy receptor function (Aoki *et al*., 2011).

A missing link in our understanding of the phosphorylation-mediated regulation of mitophagic trafficking has been the identity of the opposing phosphatase which counteracts the function of pkp1/2 on reporter proteins such as Mdh1-GFP. In the regulation of PDC catalytic activity, it is aup1/Ptc6 which was shown to dephosphorylate S313 on pda1 (Guo *et al*., 2017a). However the genetic interactions which we uncovered between aup1 and pkp2 make it impossible for aup1 to be playing a similar role in the regulation of mitophagic trafficking. The data shown in Fig. 4D and 4E allows us to suggest Ptc5 as a candidate for a phosphatase which opposes the phosphorylation of matrix proteins by pkp1/2. While further work will be required in order to determine whether this effect is direct, this result further differentiates the PDC regulatory interactions impinging on pyruvate dehydrogenase activity, from the signaling cascade which regulates the mitophagic trafficking of individual protein species.

### Ideas and speculations

A central result of this study is the discovery of a role for the PDC in regulating matrix protein phosphorylation, independent of its catalytic activity. This is underscored by the fact that the K75R mutation in the E2 subunit Lat1, a mutant which cannot be conjugated to lipoic acid- and is therefore a loss of function mutant (Lawson *et al*., 1991)-actually shows significantly higher levels of phosphorylation and mitophagic trafficking of Mdh1-GFP (see Figures 2G and 4F). Structurally, the *lat1*^K75R^ mutation is of great interest, because it forces the PDC into a configuration where most or all E1 subunits carry acetylated TPP, due to a lack of an efficient thiol acceptor. While this may be an unusual combination under normal physiological conditions, it would be expected under conditions where NAD^+^ (or FAD) is limiting, because a lack of NAD^+^ would prevent regeneration of oxidized lipoic acid. We therefore suggest that this result supports a model where the PDC can sense metabolite concentrations to direct downstream phosphorylation events.

We propose, based on our data, that the PDC can function as a signaling hub which regulates protein phosphorylation in the mitochondrial matrix in response to metabolic cues (see model in Figure 8). We hypothesize that allosteric transitions within the PDC, induced by ligand or substrate binding, will affect the structure of the complex, leading to changes in the activity of its associated kinases and phosphatase(s). The entire system, including the redundancy of the pyruvate dehydrogenase kinases (there are 4 PDKs in mammals) and the attendant PDC phosphatase, seems to be highly conserved. Strikingly, the region around the yeast Mdh1 phosphorylation site at Thr199 (IGGHSGITIIPPLISQ), which we previously showed to be crucial for mitophagic trafficking, is also highly conserved in human MDH2 (IGGHAGKTIIPLISQ) and the homologous Thr204 in the human protein was also reported to be phosphorylated (Kettenbach *et al*., 2011). Therefore, it is likely that these reciprocal regulatory interactions are also conserved. Indeed, elegant biochemical studies from the Roche lab demonstrated that the activation of human PDK2 by NADH and acetyl coA was due to bound E2 becoming acetylated in the partial reverse reaction, demonstrating the existence of this exact type of allosteric coupling in human PDKs (Baker *et al*., 2000), although in that case, their conclusions limited the implication to the allosteric regulation of E1α phosphorylation. Our results now suggest a different interpretation for these previous findings, wherein allosteric shifts in the E2 may regulate the phosphorylation of additional substrates. Similarly, it was recently reported that the overexpression of *PDK4* enhances mitophagy in mammalian cells (Park *et al*., 2015)· although again, the assumption was that this reflects regulation of pyruvate dehydrogenase activity. Another recent study found that branched-chain amino acids allosterically modulate PDC catalytic activity (Li *et al*., 2017), supporting the notion that conformational changes in the PDC occur in response to metabolic cues. The results we have presented here suggest an additional dimension to the picture, by indicating that the PDC functions upstream of the PDKs to regulate mitochondrial function in response to such conformational changes.

**Figure 8.**
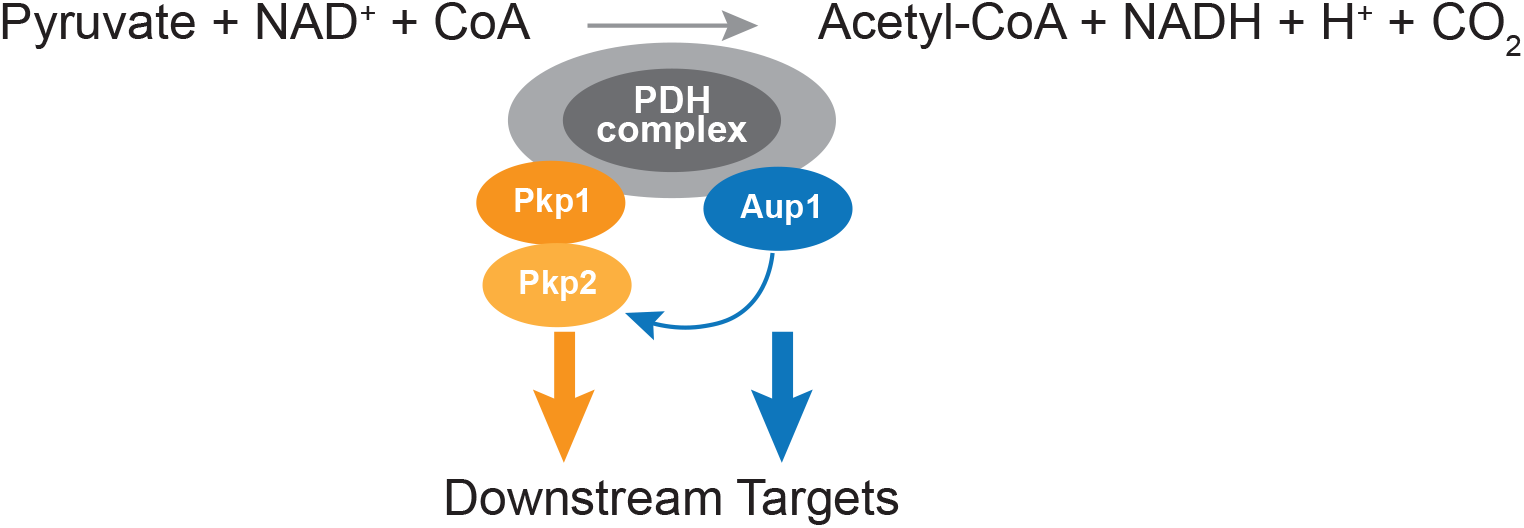
Model for regulation of mitochondrial protein phosphorylation by the PDC. We propose that the known stable physical interactions between aup1 and pda1 (ref. 22) and between pkp1 and pda1 (ref. 23) can function as an allosteric interface to regulate aup1, pkp1 and pkp2 activity towards other proteins within the mitochondrial matrix. We previously showed that pkp2 functions downstream of aup1, and in this study we showed that cooverexpression of pkp1 and pkp2 completely overcomes the mitophagy defect in *pda1Δ* cells. Since we also demonstrate that the mitophagy-related function of aup1 is independent of pda1 phosphorylation, we propose that pda1-bound aup1 functions downstream of pda1 to transduce signals to the kinase(s) in the pathway. It remains to be seen whether physiological PDC ligands can allosterically regulate phosphorylation in the matrix.

The concept of large signaling complexes that integrate metabolic cues to regulate cellular activity has been the focus of intense study in recent years. TORC1 and TORC2 are example of such mega-complexes, which integrate nitrogen, amino acid and carbohydrate metabolism with hormone signaling, to generate readouts that regulate cellular responses to changes in the environment (Eltschinger and Loewith, 2016; Nicastro *et al*., 2017; Szwed *et al*., 2021). We hypothesize that the PDC could serve a similar function within mitochondria, by regulating matrix protein phosphorylation in response to changes in metabolite levels. Further work will now be required to test specific aspects of this hypothesis.

## Methods

### Yeast strains, plasmids and growth conditions

Yeast strains used in this study are listed in Supplementary Table I. Deletion mutants and epitope-tagged strains were generated by the cassette integration method (Longtine *et al*., 1998). Strain HAY75 (MATα *leu2-3,112 ura3-52 his3-Δ200 trp1-Δ901 lys2-801 suc2-Δ9*) was used as the WT genetic background in this study..

All knockout strains were verified by PCR. Oligonucleotides used in this study are detailed in Supplementary Table III, and plasmids are detailed in Supplementary Table II.

Yeast were grown in synthetic dextrose medium (0.67% yeast nitrogen base w/o amino acids (Difco), 2% glucose, auxotrophic requirements and vitamins as required) or in SL medium (0.67% yeast nitrogen base w/o amino acids (Difco), 2% lactate pH 6, 0. 1% glucose, auxotrophic requirements and vitamins as required). All culture growth and manipulation were at 26 ^0^C. Yeast transformation was according to Gietz and Woods (Gietz and Woods, 2002).

For nitrogen starvation experiments, cells were grown to mid-log phase (OD_600_ of 0.4-0.6) in synthetic dextrose medium (SD), washed with distilled water, and resuspended in nitrogen starvation medium (0.17% yeast nitrogen base lacking amino acids and ammonium sulfate (Difco), 2% glucose) for the times indicated in individual experiments. Overexpression studies with *CUP1* promoter-based vectors were carried out by supplementing the medium with 5 μM CuSO_4_, for both control (empty vector) and overexpressing cells.

### Chemicals and antisera

Chemicals were purchased from Sigma-Aldrich (Rehovot, Israel) unless otherwise stated. Custom oligonucleotides were from Hylabs (Rehovot, Israel). Anti-GFP antiserum was from Santa-Cruz (Dallas, TX). Horseradish peroxidase-conjugated goat anti-rabbit antibodies were from MP Biomedicals (Santa Ana, CA). Anti-phosphoserine/threonine antibodies were from ECM Biosciences (catalog # PM3801).

### Preparation of whole-cell extracts for western blot analysis

Cells (10 OD_600_ units) were treated with 10% cold trichloroacetic acid (TCA) and washed three times with cold acetone. The dry cell pellet was then resuspended in 100 μl cracking buffer (50 mM Tris pH 6.8, 6 M urea, 1 mM EDTA, 1% SDS) and vortexed in a Heidolf Multi Reax microtube mixer at maximum speed with an equal volume of acid-washed glass beads (425-600μm diameter), for a total of 30 min. Lysates were clarified by centrifugation at 17,000 xg for 5 min and total protein was quantified in the supernatant using the BCA protein assay (Thermo Scientific, Rockford, IL). SDS loading buffer (final concentrations of 100 mM Tris pH 6.8, 20% glycerol, 2% SDS, 500 mM β-mercaptoethanol) was added to the lysate and the samples were warmed to 60 °C for 5 min prior to loading on gels. ImageJ software was used for band quantification.

### Free GFP release assays for analyzing mitophagic trafficking efficiency

Quantitative analysis of mitophagic trafficking was analyzed using GFP chimeras of endogenous mitochondrial matrix proteins or an artificial mitochondrially-targeted GFP chimera of mouse cytoplasmic DHFR. This assay relies on the resistance of GFP to proteolytic degradation in the yeast vacuole relative to that of other proteins. Since GFP molecules derived from different chimeras can be expected to have the same lifetime, this allows a semi-quantitative analysis of the efficiency of mitophagic targeting as measured by the percentage of the total signal that is converted to free GFP (Klionsky *et al*., 2016; Kolitsida and Abeliovich, 2019).

### Immunoblotting

SDS-10% polyacrylamide gels were transferred to nitrocellulose membrane by using either a wet or semidry blotting transfer system. Membranes were blocked for 1 h with TPBS (137 mM NaCl, 2.7 mM KCL, 10 mM Na_2_HPO_4_, 1.8 mM KH_2_PO_4_, 1 % Tween and 5 % milk powder (BD Difco Skim Milk), followed by incubation with rabbit anti-GFP (1:5000) for 1.5 h, then washed 4x 2 min with TPBS, incubated for 1.5 h with HRP conjugated secondary antibody (1:10000). The membranes were then washed 4x with TPBS, incubated with SuperSignal Chemiluminescence substrate (Thermo Scientific), and exposed to a Molecular Imager ChemiDoc^TM^ XRS imaging system. Where necessary, 2% normal goat serum was added to blocking solutions to reduce background.

### Calculation of statistical significance of differences in % free GFP signals from immunoblots

Data (% free GFP) were log-transformed before analysis in order to stabilize variances. A repeated measures ANOVA was performed to compare mutants and proteins simultaneously. Thereafter, mutants were compared for each protein and proteins were compared for each mutant by the Tukey HSD test (p<0.05). ANOVA analysis was carried out using JMP 12 software.

### Native immunoprecipitation

Cells (20 OD_600_ units) were washed 3x with 1 ml of cold 10 mM PIPES pH 7, 2 mM PMSF. The cell pellet was washed in three volumes of cold lysis buffer (20 mM HEPES pH 7.4, 50 mM NaCl, 1mM EDTA, 10% glycerol, 0.2 mM Na_3_VO_4_, 10 mM H_2_NaPO_4_, 10 mM β-glycerophosphate, 10 mM NaF, 10% phosphate cocktail inhibitors, 1 mM PMSF, proteases inhibitors (final concentrations: 5 μg/ ml antipain, 1 μg/ml aprotinin, 0.5 μg/ml leupeptin, 0.7 μg/ml pepstatin). The cells were then frozen in liquid nitrogen and stored at −80°C. Extracts were prepared from the frozen pellets using a SPEX Freezer/Mill according to the manufacturer’s instruction. The frozen ground yeast powder was thawed at room temperature, and equal amount of lysis buffer plus Triton-X100 was added to a final concentration of 0.5% v/v detergent. Cell debris was removed by centrifugation at 17.000x g for 10 min at 4°C. 20 μl of GFP-Trap bead slurry (Chromotek) was pre-equilibrated in lysis buffer, per the manufacturer’s instructions. For co-immunoprecipitation experiments (Figure 6), agarose-conjugated anti-HA antibody (Santa Cruz) was used instead of GFP-trap. The cleared lysate was added to the equilibrated beads and incubated for 1 h at 4 °C with constant mixing. The beads were washed five times with 1 ml ice cold wash buffer (20 mM HEPES pH 7.4, 50 mM NaCl, 0.5 mM EDTA, 10% glycerol, 0.5% Triton-X, 0.2 mM NaOV_3_, 10 mM H_2_NaPO_3_, 10 mM β-glycerophosphate, 10 mM NaF, resuspended in 50 μl 2x SDS-loading buffer (200 mM Tris pH 6.8, 40% glycerol, 4% SDS, 1 M β-mercaptoethanol) and boiled for 10 min at 95°C. For phosphatase treatment, immunoprecipitated beads were incubated with 40 Units of lambda protein phosphate (New England Biolabs) in a 50 μl reaction. Samples were washed once with wash buffer and resuspended in 50 μl 2 x SDS-loading buffer.

### SILAC labeling

To quantify relative phosphorylation on mitochondrial proteins in WT versus *pda1Δ* backgrounds, cell cultures of each strain were grown separately in SL medium supplemented respectively with 0.003% (w/v) ^13^C_6_ ^15^N2 L-lysine (Lys8; Cat. Number 211604102, Silantes, Munich) and 0.001% (w/v) ^13^C_6_^15^N_4_ L-arginine (Arg10; Cat. Number 201603902, Silantes, Munich) or with identical concentrations of unlabeled lysine and arginine, in addition to the standard auxotrophy supplementation. At the indicated time point, 50 OD_600_ units of cells from each labeling regime were combined, washed with cold lysis buffer, and cells were frozen in liquid nitrogen. To lyse the cells, the frozen suspensions were processed in a SPEX Freezer/ Mill according to the manufacturers’ instructions. The frozen ground powder was thawed and clarified by centrifuging for 10 min at 500 xg. The clarified lysates were then further centrifuged for 10 min at 13,000 xg, the supernatant was discarded and the pellet was precipitated with 10% ice cold TCA and washed 3x with cold acetone.

### MS analysis

Mass spectrometric measurements were performed on a Q Exactive HF-X mass spectrometer coupled to an EasyLC 1200 (Thermo Fisher Scientific, Bremen, Germany). Prior to analysis phosphopeptides were enriched on an Automated Liquid Handling Platform (AssayMap Bravo, Agilent) using Fe (III)-NTA cartridges (5 *μ*L) essentially as described previously (PMID: 28107008). The MS raw data files were analysed by the MaxQuant software (Cox and Mann, 2008) version 1.4.1.2, using a Uniprot *Saccharomyces cerevisiae* database from March 2016 containing common contaminants such as keratins and enzymes used for in-gel digestion.

### Statistical analysis

Phosphosite SILAC ratios were normalized to respective protein changes to discriminate regulated sites from regulated proteins. Measurements of three biological replicates (Pearson correlation of 0.64-071) comparing WT to *pda1Δ* cells were averaged and log2 transformed. To determine sites that changed significantly between genotypes we calculated an outlier significance score on the merged log2-transformed phosphosite ratios according to (Cox and Mann, 2008) (significance A, p<0.05, BH corrected).

For statistical analysis of western blotting quantifications in Figures 3B, 4B, and 4G, % normalized phospho-AA signal intensity relative to GFP signal values were calculated from raw files. MANOVA was carried out for overall comparison of all the genotypes measured on three days (day 1, day 3 and day 5, N=3). Log-transformation of the data was applied before analysis to stabilize variances. Pairwise comparison of genotypes was carried out by contrast tests.

### Site directed mutagenesis

Site directed mutagenesis protocol was carried out according to the Stratagene QuikChange system. *Pfu* DNA Polymerase (Fermentas) was used with a thermocycling protocol followed by removal of the parental strands with *DpnI* digestion. Thermocycling conditions were as follows: Initial denaturation at 95°C for 2 min, then (steps 2-4), 30 sec at 95 °C, 60 sec at 55 °C, and 12 min at 68 °C. Steps 2-4 repeated for 18 cycles. PCR products were treated with DpnI (fermentas) for 1 hr at 37 °C, and precipitated using with 70% ethanol. DNA pellets were then dried and resuspended with 20 μl TE buffer pH 8. All mutagenesis products were transformed into *E. coli* DH5α and mutations were verified by sequencing. The oligonucleotide primers used to create site-directed MDH1-GFP variants and combined variants are listed in Supplementary Table III.

### Fluorescence microscopy

Culture samples (3 μl) were placed on standard microscope slides and viewed using a Nikon Eclipse E600 fluorescence microscope equipped with the appropriate filters and a Photometrics Coolsnap HQ CCD. To achieve statistically significant numbers of cells per viewing field in the pictures, 1 ml cell culture was collected and centrifuged for 1 min at 3,500 xg prior to sampling. For quantitative analysis of co-localization, ImageJ software with the colocalization Plugin, JACoP (Bolte and Cordelières, 2006) was used. Representative fields were analyzed in terms of channel intensity correlation coefficient.

For confocal microscopy, cells were placed on standard microscope slides and micrographs were obtained by confocal laser scanning microscopy using a Leica SP8, using a 63x water immersion lens. GFP excitation was carried out using the 488 nm line, and emission was collected between 500-540 nm. For RFP excitation we used the 552 nm line and fluorescence emission was collected between 590-640 nm. Deconvolution of Z-stacks was carried out using ImageJ software.

For statistical analysis of channel overlap data, ANOVA analysis was carried out using JMP 16.0 software. Correlations were compared over mutations and days by two-factor ANOVA. As the mutation X day interaction effect was statistically significant (p<0.0001), the two days were compared for each mutation by contrast t-tests. Pre-planned comparisons to WT were also performed by contrast t-tests.

#### Determination of colony forming efficiency

Cells (HAY75 and PKY832), harboring empty vectors or plasmids overexpressing pkp1 and pkp2, were grown to late log phase in SD medium, washed and equal cell numbers were resuspended and incubated in SL medium for 5 days. After 5 days, the cells were re-counted, diluted to an expected confluency of 175 colonies/plate, and plated on YPD plates. See Supplementary Figure 5 for an example.

## Acknowledgements

We wish to thank J.C. Martinou and Maya Schuldiner for discussions, Hillary Voet for help with statistical analysis, and Einat Zelinger for confocal microscopy. This work was funded by ISF grant 445/17 (to HA), GIF grant I-111-412.7-2014 (to HA and JD), the University and Canton of Fribourg and the Swiss National Science Foundation (J.D.).

## Author contributions

P.K. planned experiments, carried out experiments, generated figures, and wrote the manuscript. V. N. carried out experiments. J.Z. carried out MS analyses. N.N. planned experiments and wrote the manuscript. J.D. carried out MS analyses, planned experiments, generated figures, and wrote the manuscript. H.A. Planned experiments, generated figures, and wrote the manuscript.

## Supplementary Figure Legends

**Supplementary Figure 1.** Deletion of *PDA1* does not affect the levels of pkp1, pkp2, or aup1. Cells expressing WT pda1 (WT) or *pda1Δ* cells expressing pkp1-HA, pkp2-HA or aup1-HA were grown in SL as described in the legend to Figure 1, and samples were taken after 1 day of incubation. protein extracts were analyzed by anti-HA immunoblotting to compare the level of the HA tagged protein in WT versus *pda1Δ* extracts. Untagged control extract was probed in parallel in order to ascertain the identity of the immunoreactive band. The data shown is representative of 3 independent biological experiments.

**Supplementary Figure 2**. **Mutations in *LAT1* do not affect mitophagic trafficking of mtDHFR-GFP.** *lat1Δ* cells (PKY2077) harboring empty vector or plasmids expressing Lat1 and Lat1K75R, and expressing mtDHFR-GFP, were grown in SL medium for the indicated amounts of time and protein extracts were generated. The extracts were analyzed by immunoblotting with *α*-GFP antibodies. The results represent an average of at least 3 independent biological repeats.

**Supplementary Figure 3**. Deletion of *pda1* causes hypophosphorylation of Qcr2-GFP. WT (PKY2255) and *pda1Δ* (PKYPKY2256) cells expressing Qcr2-GFP were grown in SL medium for the indicated amounts of time and protein extracts were generated and immunoprecipitated with *α*-GFP antibody. The immunoprecipitates were then probed with *α*-phosphoamino acid antibody (top panel) and *α*-GFP antibody (bottom panel). Right panel: day 5 samples were treated overnight with lambda phosphatase or mock treated as described in “Methods”, and analyzed by immunoblotting.

**Supplementary Figure 4**. Deletion of *pda1* causes hypophosphorylation of Aco1-GFP. WT (PKY2257) and *pda1Δ* (PKY2258) cells expressing Aco1-GFP were grown in SL medium for the indicated amounts of time and protein extracts were generated and immunoprecipitated with *α*-GFP antibody. The immunoprecipitates were then probed with *α*-phosphoamino acid antibody (top panel) and *α*-GFP antibody (bottom panel). Right panel: day 5 samples were treated overnight with lambda phosphatase or mock treated as described in “Methods”, and analyzed by immunoblotting.

**Supplementary Figure 5**. Illustration of the effects of *pda1* deletion on colony formation efficiency. Cells (HAY75 and PKY832), harboring empty vectors or plasmids overexpressing pkp1 and pkp2, were grown to late log phase in SD medium, washed and equal cell numbers were resuspended and incubated in SL medium for 5 days. After 5 days, the cells were recounted, diluted to an expected confluency of 175 colonies/plate, and plated on YPD plates. WT (left picture) cells gave rise to 103 colonies (59%), *pda1Δ* cells (middle picture) gave rise to 44 colonies (25%) and co-overexpression of pkp1 and pkp2 plasmids (right picture) in *pda1Δ* cells gave rise to 98 colonies (56%). Three such experiments were used to generate Figure 4C.

