## Supplemental Figures for "The pyruvate dehydrogenase complex regulates matrix protein phosphorylation and mitophagic selectivity"

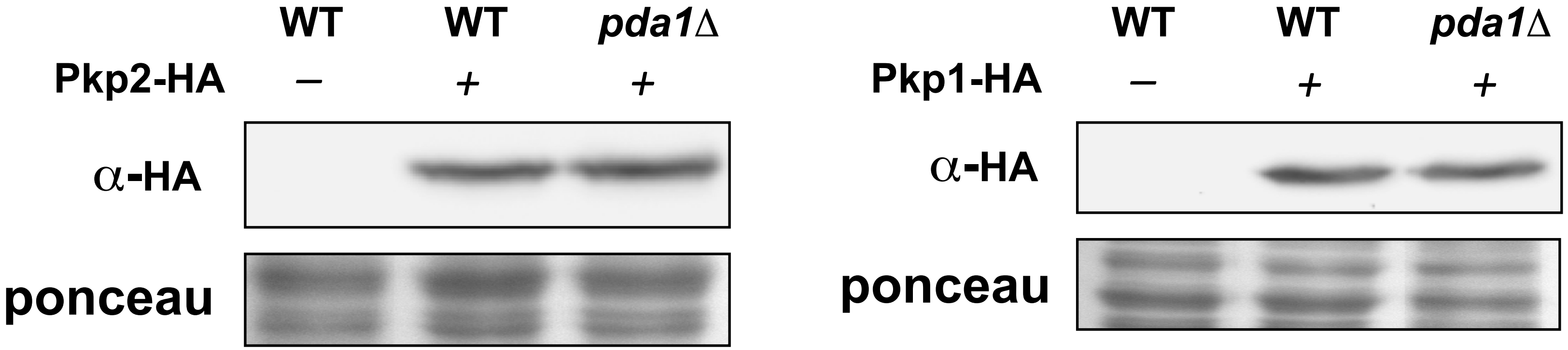


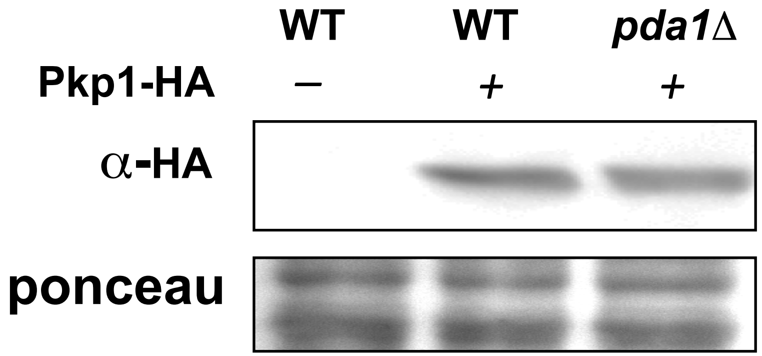


**Supplementary Figure 1**

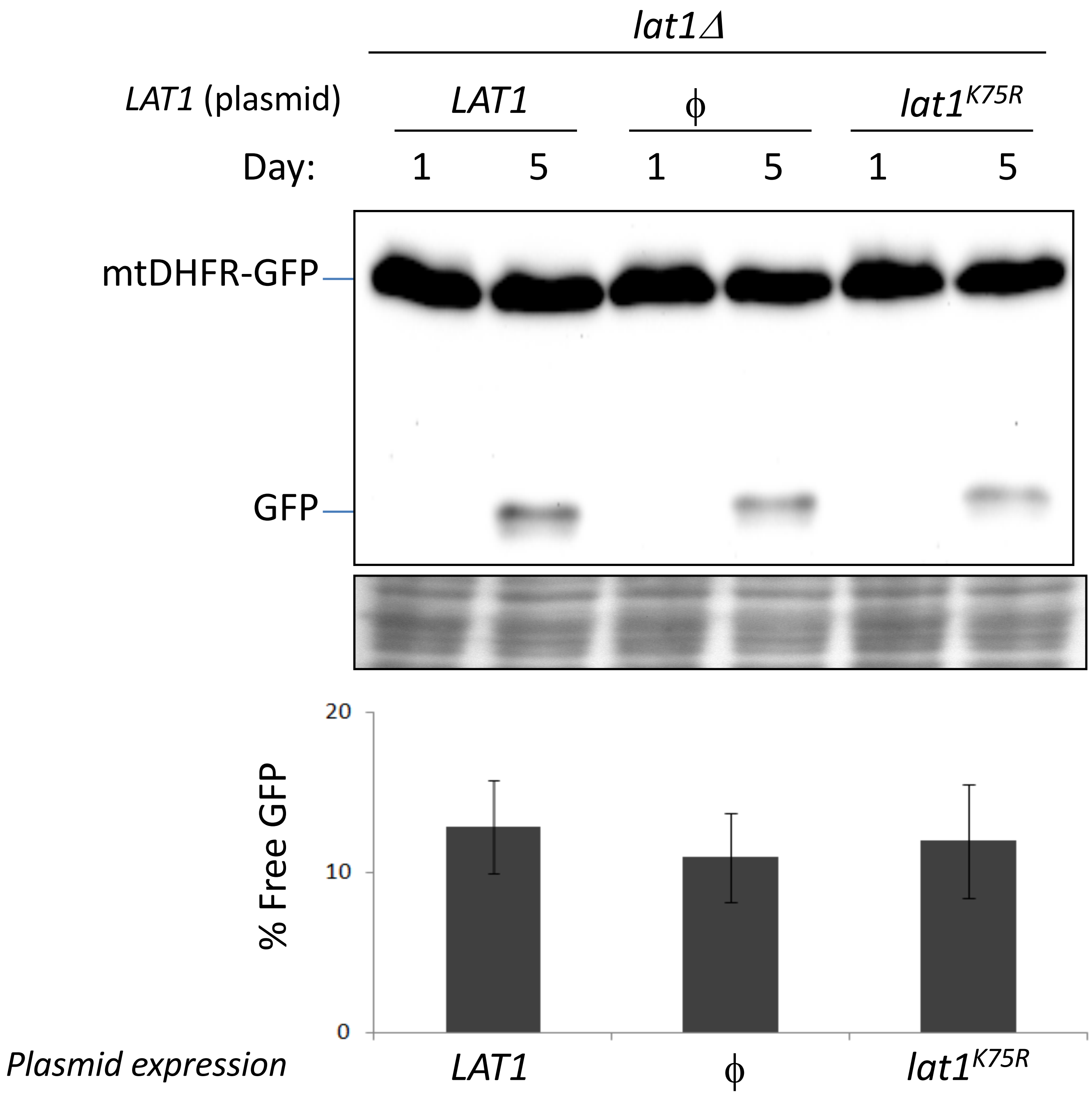


Supplementary Figure 2

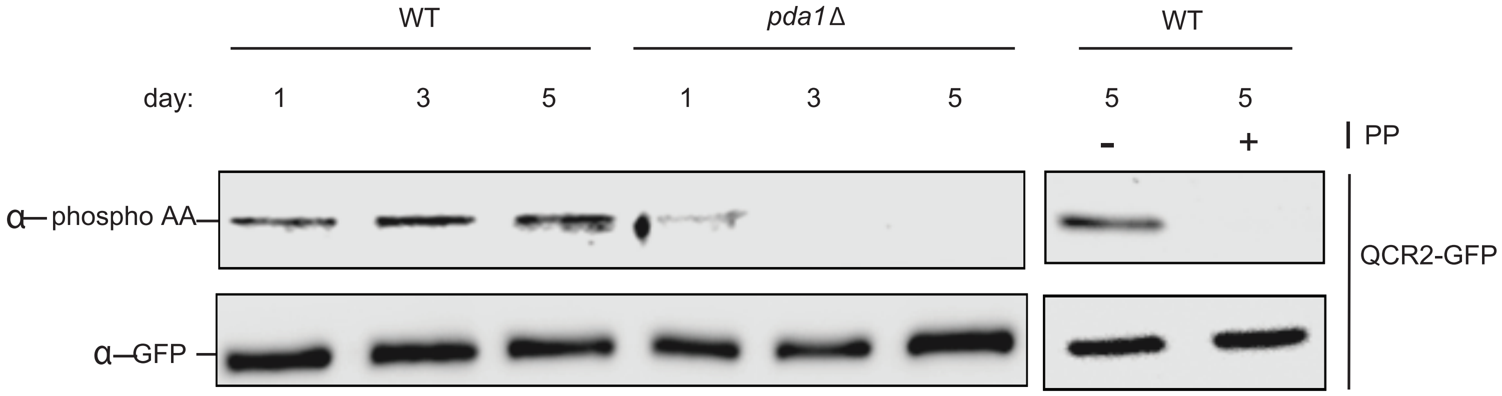


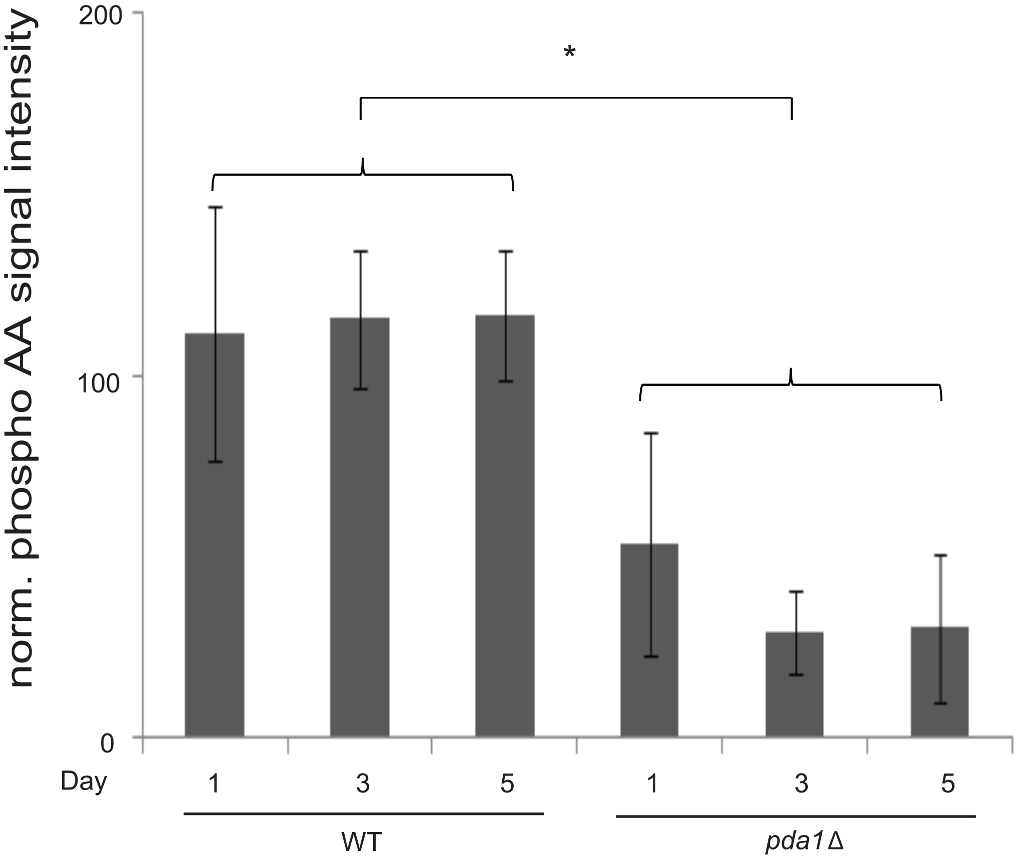


Supplementary Figure 3

A

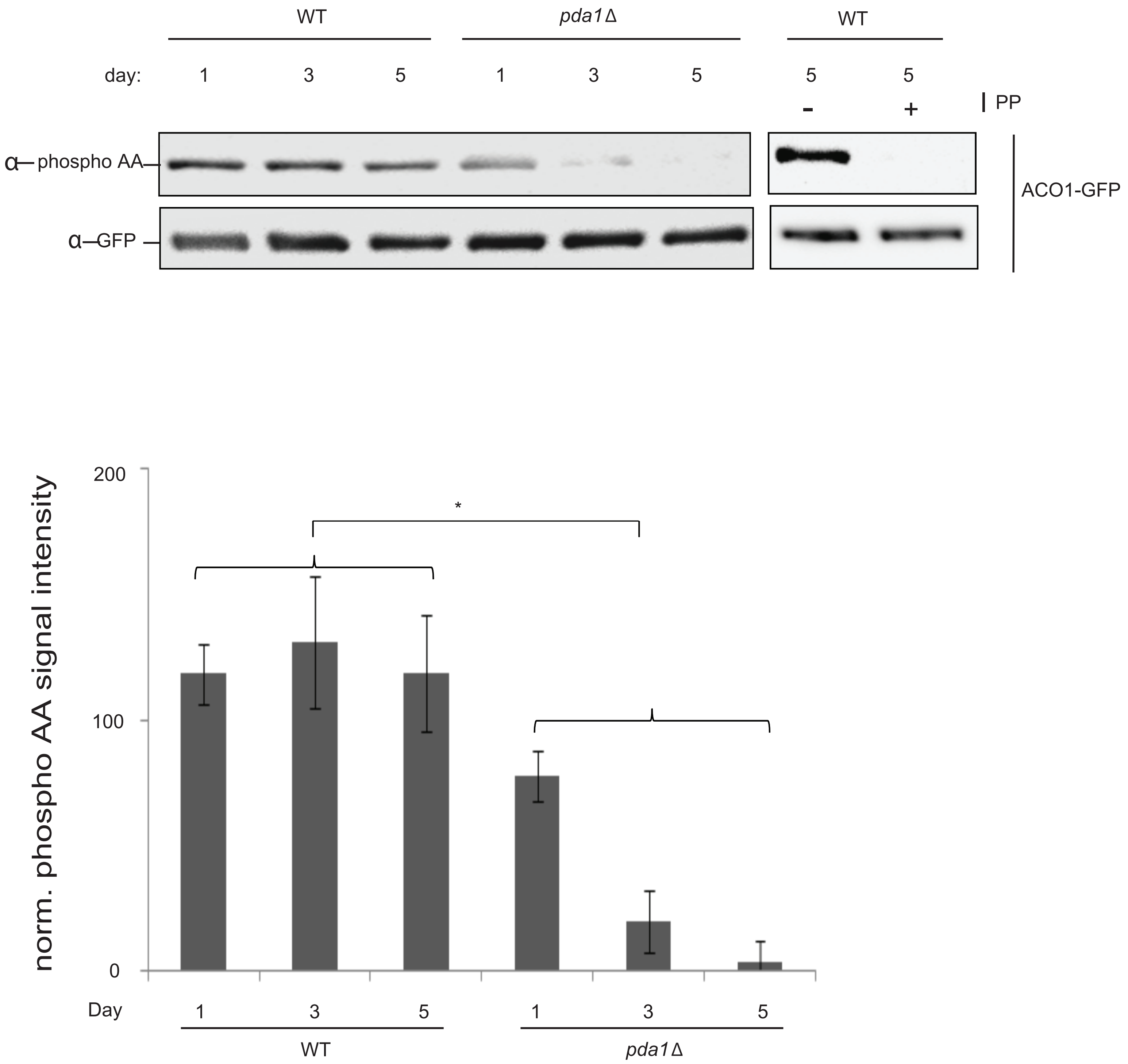


Supplementary Figure 4

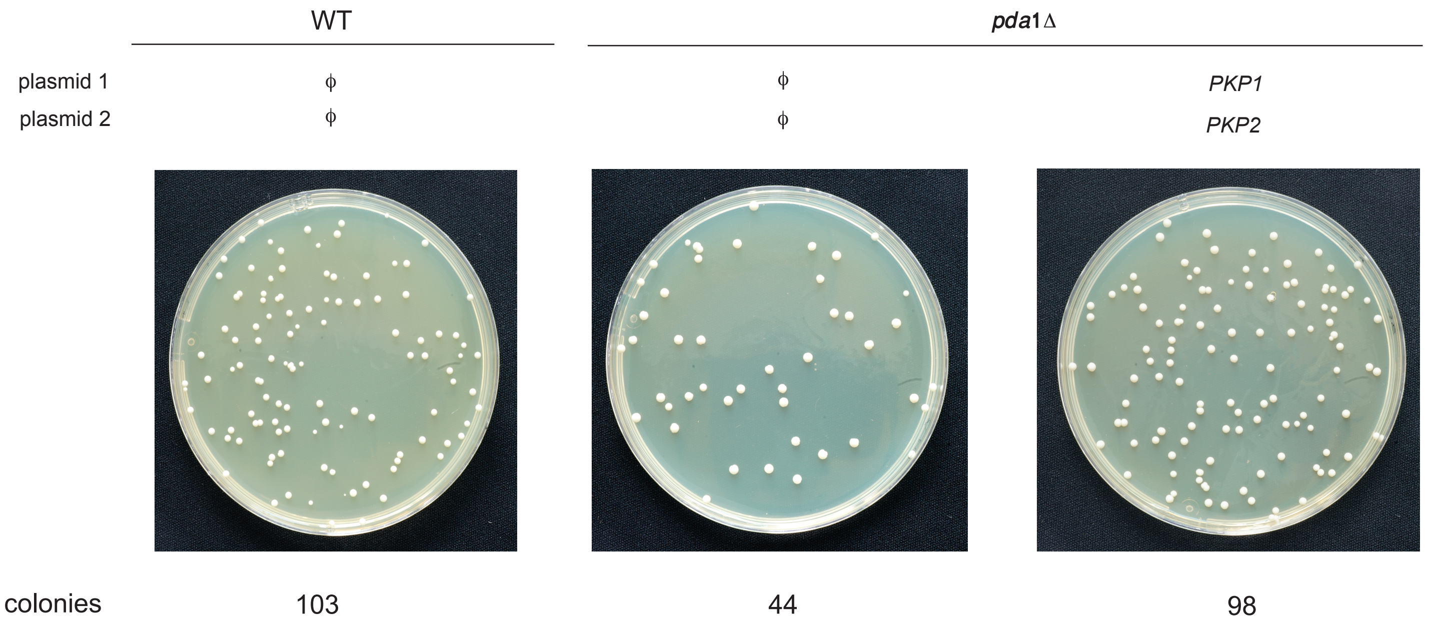


Supplementary Figure 5
